# VFB-MCP: Natural-Language Access to *Drosophila* Neuroscience Grounded by an Expert-Curated Ontology-Led Knowledgebase

**DOI:** 10.64898/2026.06.16.732577

**Authors:** Alex D McLachlan, Robert Court, Clare Pilgrim, Kit Longden, Nicolas H Brown, David Osumi-Sutherland, Gregory S X E Jefferis, J Douglas Armstrong

## Abstract

Biological databases store curated knowledge that researchers traditionally access through web interfaces or APIs. To move beyond casual browsing requires domain-specific knowledge and expertise to frame the queries necessary to explore this data. This generates barriers to both experienced users trying to integrate wide-ranging sources of information and for new users entering scientific fields undergoing paradigm shifts such as connectomics. A potentially powerful method to lower these barriers is to facilitate natural-language access to these databases by exposing them to large language models (LLMs) via the Model Context Protocol (MCP). Here we implement this for Virtual Fly Brain (VFB), an expert-curated and ontology-backed knowledgebase of *Drosophila* neuroscience, providing the precision needed to make recently integrated connectome and molecular data accessible. Benchmarked on 30 neuroscience tasks against a bare LLM and a web-search-assisted LLM, the VFB-MCP-equipped LLM produces precise, verifiable and appropriately quantified answers on 25/30 tasks vs 14/30 for web and 2/30 for bare. The MCP advantage is largest for tasks where data quantification is required (89% vs 11% web). This work establishes MCP layered over ontology-backed knowledge graphs as an effective method to improve LLM response quality for neuroscience and connectomics data, promoting accessibility and accelerating the reliable analysis of complex integrated datasets.

## Introduction

Expert curation has produced large, structured knowledgebases which formalise knowledge from fields across the life sciences, such as intra- and intercellular signalling (OmniPath) and biological pathways (Reactome).^1,2^ Integrated knowledgebases are also emerging such as the Alliance of Genome Resources (Alliance) which is a coalition of model organism knowledgebases covering eight species with ongoing harmonisation between them.^3^ Many of these resources are formalised through domain-specific ontologies such as the Gene Ontology,^4^ Drosophila Anatomy Ontology (DAO),^5^ or the multi-species anatomy ontology Uberon,^6^ and stored in graph databases, where entities and their typed relationships form traversable networks ^7^. Yet an accessibility gap separates these extensive and highly formalised resources from researchers, and this is particularly acute when they have newly entered a field. Web interfaces support browsing but become cumbersome for complex queries. Programmatic APIs provide deep flexibility but require coding expertise. In both cases intimate knowledge of the underlying data models is required to deeply query and understand the data returned. Further, research often requires multiple query steps and integration of the results, for which reasoning is a requirement unmet by existing access methods.

Large language models (LLMs) present a third interface for these knowledgebases; natural-language.^8^ The dominant approach for grounding LLMs in external knowledge is retrieval-augmented generation (RAG), which retrieves relevant external information and includes it in the LLM’s context window to inform completion.^9^ RAG has been applied to biological and biomedical questions with promising results: genetic variant annotations,^10^ synthetic biology,^11^ and general biology.^12^ However, traditional RAG pipelines rely on static, bespoke retrieval. The pipeline retrieval step occurs before the LLM receives the prompt, and therefore the LLM has no agency in determining what context is required and provided. The LLM must then extract answers from this context, leading to well-documented problems including imprecise retrieval, confabulated citations and the ‘lost-in-the-middle’ phenomenon where key information is overlooked in long contexts.^13^ Importantly, performance also depends on the interaction between the context provided and the type of question asked. For example, RAG-enabled LLMs performed worse on neurology case-based questions compared to knowledge-inquiring questions, possibly due to the specificity of the case-based questions compared to the context given.^14^

The Model Context Protocol (MCP) takes a different approach.^15^ Rather than retrieving context upstream of the LLM, MCP defines a standardised interface through which LLMs invoke tools at runtime: structured functions with typed inputs that query databases, execute analyses and return results directly into the completion. The LLM acts as an agent that reasons about which tools to call, in what order, and how to combine their outputs. Importantly, the results of one tool call inform the next, allowing multi-hop retrieval. MCP therefore implements the agentic RAG paradigm,^16^ but through a standardised protocol enabling dynamic multi-hop retrieval without bespoke tool integration. Further, MCP is model-agnostic, allowing any model that supports MCP to be used. A growing set of biological MCP servers has begun to exploit this paradigm, wrapping REST APIs for bioinformatics web services (MCPmed)^17^, exposing GraphQL APIs for clinical knowledgebases (CIViC MCP) ^18^, and auto-generating tools for biomedical knowledge graphs (Samyama).^19^

To show that augmenting LLMs with biological database tools via MCP is useful, the quality of the answers given must be benchmarked. Where MCP servers exist they are often provided as-is to researchers with no benchmarking of performance: MCPmed,^17^ Alliance-MCP,^20^ EBI Ontology Lookup Service (OLS).^21^ However, MARRVEL-MCP provides 44 tools for Mendelian-disease variant interpretation and reports a 94% pass rate on a 100 question task battery versus 41% without MARRVEL-MCP with the lightweight model gpt-oss-20b.^22^ Samyama reports 98% accuracy with MCP tools vs 85% with schema-aware text-to-Cypher and 75% bare on 40 pharmacology questions (automated review).^19^ These results establish empirically that an MCP layer over a curated biological resource lifts LLM accuracy substantially.

We demonstrate this approach in the domain of *Drosophila* neuroscience, where the accessibility challenge is acute: new connectomes catalogue hundreds of thousands of neurons and tens of millions of synapses with larval^23^ and adult^24–27^ brain examples available. At the same time single-cell transcriptomic atlases^28^ and transgenic driver collections^29,30^ provide complementary views of neural-circuit organisation. Virtual Fly Brain (VFB; https://virtualflybrain.org) integrates these resources for *Drosophila* under a shared semantic framework built on the DAO, stored in a Neo4j graph database.^31^ Here we present VFB-MCP, a server that exposes this graph-structured knowledge as a set of composable tools and is, to our knowledge, the first MCP implementation for neuroscience and connectomics data. We benchmark this system on a graded 30-task battery against bare-LLM and web-search alternatives, with expert review and using a current frontier LLM (Opus 4.7), and quantify both the gains and the specific failure modes that the architectural design prevents.

## Results

### Architecture: a small generic MCP surface with query introspection

A core advantage of allowing an LLM to query the VFB knowledgebase (kb) directly is giving the LLM the ability to decide what context it needs. Therefore, we designed the MCP tools with a simple architecture that allows sequential context collection and introspection at runtime (Figure 1A). A set of three core, composable tools for querying the kb is presented with a MCP preamble that informs the LLM of when to request each one, and what it does. Only using relevant context helps to minimises token usage and manages context window use. The architecture relies on three layers: a formal ontology that defines the types, relationships, and constraints of the domain; a graph database that stores curated knowledge as typed nodes and edges conforming to the ontology; and a minimal MCP tool surface that exposes search, entity introspection, and structured-query execution.

**FIGURE 1:**
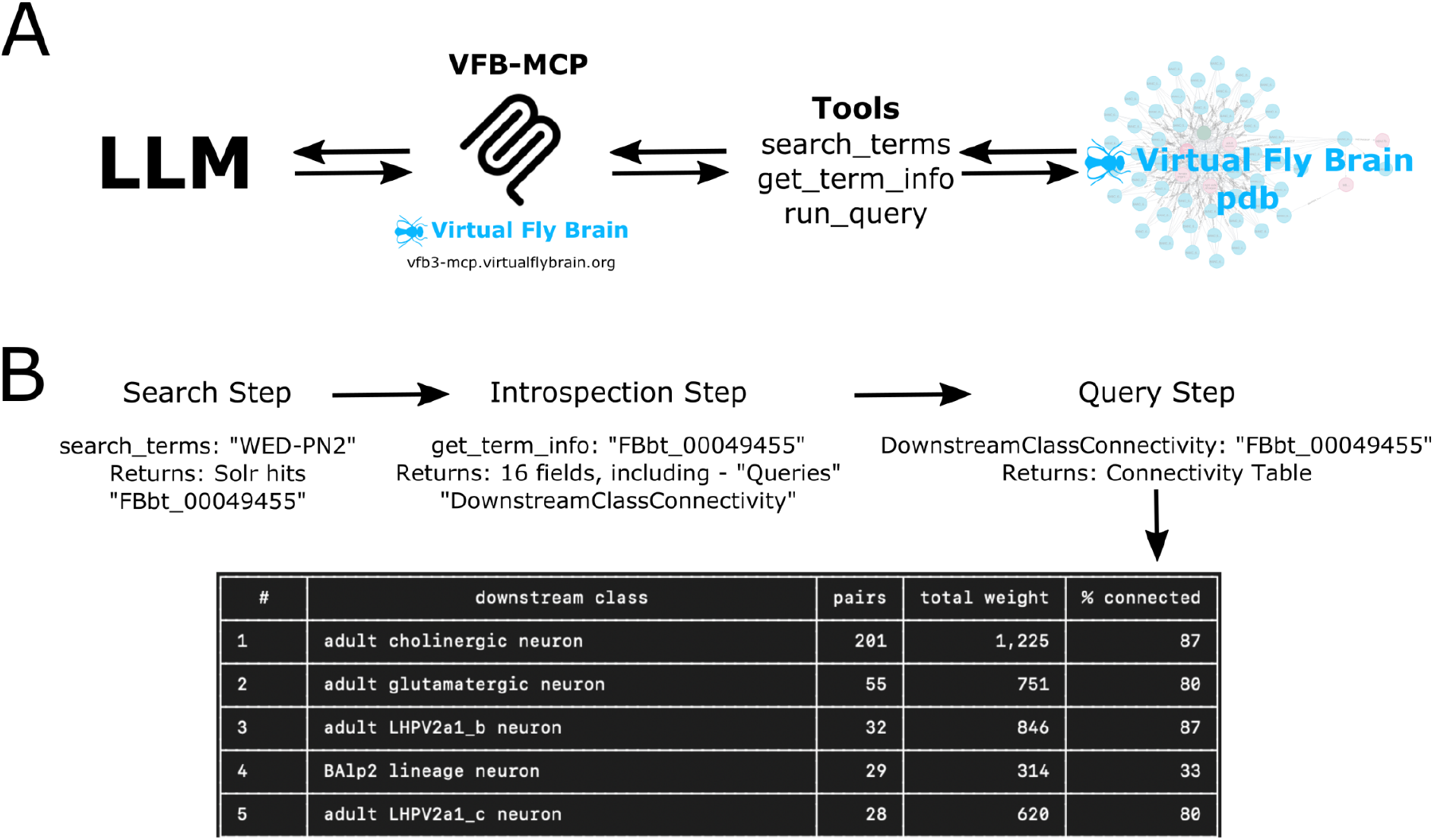
Architecture and the ontology-backed MCP pattern. **(A) VFB-MCP implementation:** LLM client → MCP protocol → VFB-MCP server (3 generic tools) → Neo4j graph DB / Solr index. The per-entity `Queries` array (returned by `get_term_info`) carries the valid query types for that entity, enabling a small tool surface to cover the breadth of biological questions the system is asked. **(B) Example of tool use flow:** search step > introspection step > query step and response from the MCP server query tool call.

We implement this architecture as VFB-MCP, a TypeScript server exposing Virtual Fly Brain’s Neo4j graph database through **three generic tools**:

1. **search_terms**: Solr-backed full-text search across the VFB ontology, filterable by ~100 entity-type facets (anatomy, neuron classes, neurotransmitter classes, lineages, datasets, etc.). Returns the matching VFB ID(s) with metadata.
2. **get_term_info**: fetches the canonical record for one or more VFB IDs. The returned record includes a `Queries` array enumerating the **valid** `query_type` **values for that specific entity**, facilitating runtime introspection.
3. **run_query**: executes a (`id, query_type`) pair, with batch support for both shared and mixed combinations. Common query types include `PaintedDomains, SubclassesOf, NeuronInputsTo, NeuronNeuronConnectivityQuery, SimilarMorphologyTo, ListAllAvailableImages, ExpressionOverlapsHere, clusterExpression.`

Each tool call returns structured data (typed entities with identifiers, quantitative connectivity weights, hierarchical tree structures) that the LLM can reason over precisely. The graph database backend means relationship traversals (connectivity paths, ontological hierarchies, part-whole decompositions) are performed by the queries provided rather than text-extraction tasks for the LLM. The server is deployed as a stateless HTTP endpoint (https://vfb3-mcp.virtualflybrain.org) compatible with any MCP client, including Claude Desktop, Claude Code, GitHub Copilot, VS Code, and custom applications. Configuration requires a single JSON entry specifying the server URL.

The workflow for standard MCP tool use is therefore: The LLM receives the user prompt, identifies the entities of interest and requests `search_terms` to retrieve the matching VFB IDs. In the next turn, these IDs are fed back into the LLM prompt and used to request `get_term_info` which provides the possible queries. The queries are again fed back to the LLM prompt and used to request `run_query` with the appropriate query. The query results are then reasoned over to produce the final completion (Figure 1B).

### Benchmark: bare LLM vs LLM + VFB-MCP vs LLM + web

To evaluate VFB-MCP’s effectiveness, we designed a battery of 30 *Drosophila* neuroscience tasks across four categories which broadly reflect VFB use cases:

– **Category 1: Simple knowledge** (8 tasks): Well-established facts (e.g., major subdivisions of the mushroom body). Likely can be answered accurately from training knowledge alone.
– **Category 2: Integrative / survey** (5 tasks): Open-ended research questions regarding well-studied neurons/regions (experimental planning, data-availability surveys). Likely can be answered accurately from training knowledge alone but requires qualitative synthesis.
– **Category 3: Multi-hop** (8 tasks): 2–3 chained operations requiring information passing (e.g., neurotransmitter input mapping to a neuron class). Likely benefits from VFB-MCP access as this requires precise identification of entities in different contexts.
– **Category 4: Graph traversal / quantitative** (9 tasks): Connectivity reasoning, hierarchy navigation, cross-modal queries that require specific identifiers and synapse counts/transcriptomics data. Quantitative answers will likely require VFB-MCP access.

Each task was attempted under three conditions:

| Condition | Short name | Tool Access | Description |
| --- | --- | --- | --- |
| Bare LLM | Bare | None. | Training data recall only |
| LLM + web | Web | Web-retrieval tools. | Web search results supplemented with training data. |
| LLM + VFB-MCP | MCP | VFB kb access tools via VFB-MCP. | VFB kb query results supplemented with training data. |

We performed the comparison of the same LLM’s responses with VFB-MCP (LLM + VFB-MCP) and without (bare LLM) to quantify the impact of the VFB kb context on the quality of the resulting response. We further compared both the LLM + VFB-MCP and bare LLM to the same LLM with access to web-retrieval tools as this is the most common RAG strategy. With web tools an LLM can request papers, reviews, wikipedia articles, etc. and use this context to improve responses. This method entirely circumvents directly accessing the VFB kb. However, it is unknown how much of the VFB site has been used in LLM training data. Comparing LLM + VFB-MCP to these two alternative conditions therefore provides a benchmark against the alternative methods of using LLMs to answer *Drosophila* neuroscience questions.

Answers were scored by a VFB curator and author; Alex M^c^Lachlan, against VFB’s current database state as the primary ground truth on a 4-point scale: **−1** (fabricated; dominated by false specifics), **0** (incorrect or lacking any detail, but not caused by fabrication), **1** (generally correct, lacks specificity), **2** (precisely correct, quantified when appropriate and verifiable against current VFB data or a given reference). Two flags were recorded: **PF** (partial fabrication; answer scored 1 or 2 but contained at least one fabricated, but not essential, detail) and **LD** (literature–data discrepancy; published literature and current VFB disagree on the question). See Methods for full criteria.

### Example workflow: cross-modal query in five MCP calls

To illustrate the qualitative shape of MCP-mediated answers we present a worked category 4 task (T4.9): *“Kenyon cells receive dopaminergic input from mushroom body DANs. Based on single-cell transcriptomic data, which dopamine receptor genes do adult Kenyon cells express, and how do the expression levels compare across receptors? Where possible, distinguish γ vs α/β Kenyon cell subtypes*.*”*

The MCP-mediated answer used five distinct query steps:

1. `search_terms(“Kenyon cell”)`: identifies the Kenyon-cell parent class and subtype IDs (`FBbt_00100247` γ-KC, `FBbt_00100248` α/β-KC, `FBbt_00049828` adult-γ-KC, `FBbt_00049834` adult-α’/β’-KC).
2. `get_term_info(<KC IDs>)`: returns each entity’s metadata and `Queries` array, which includes `anatScRNAseqQuery` because the entity has the `hasScRNAseq` supertype.
3. `run_query(<KC IDs>, “anatScRNAseqQuery”)`: returns the linked scRNA-seq cluster identifiers (e.g., `FBlc0004160` for FCA Female Full γ-KC).
4. `get_term_info(<cluster IDs>)`: returns each cluster’s `Queries` array, which includes `clusterExpression`.
5. `run_query(<cluster IDs>, “clusterExpression”)`: returns the full gene-expression table per cluster (~1,000 genes each with quantitative `expression_level` and `expression_extent` fields), which the model then filters for the canonical Drosophila DA-receptor FBgns (Dop1R1 `FBgn0011582`, Dop1R2 `FBgn0266137`, Dop2R `FBgn0053517`, DopEcR `FBgn0035538`).

(note that the actual tool trace involved 43 tool rounds of which nine were VFB-MCP tool calls, 33 were infra tool calls and one was a cli tool call.)

The synthesised answer named all four canonical DA receptors with quantitative cross-subtype comparison (extent and expression level per receptor per cluster), correctly distinguishing γ vs α/β KCs. The web-mediated answer correctly named the four receptors qualitatively but could not produce cluster-level quantitation; the bare answer named the receptors from training knowledge but explicitly declined on quantitative ranking. The five-step traversal demonstrates the *search* → *introspect* → *execute* pattern. The model never needed a domain-specific “give me DA receptor expression for KCs” tool; the ontology’s per-entity query annotation guided the model from anatomical-class IDs to scRNA-cluster IDs to gene-expression tables, all via the same three generic primitives.

### LLM + VFB-MCP produced the highest proportion of quality answers

We used a frontier LLM, Claude Opus 4.7, across all 30 tasks and reviewed the results (Figure 2). The score distribution was:

**Figure 2.**
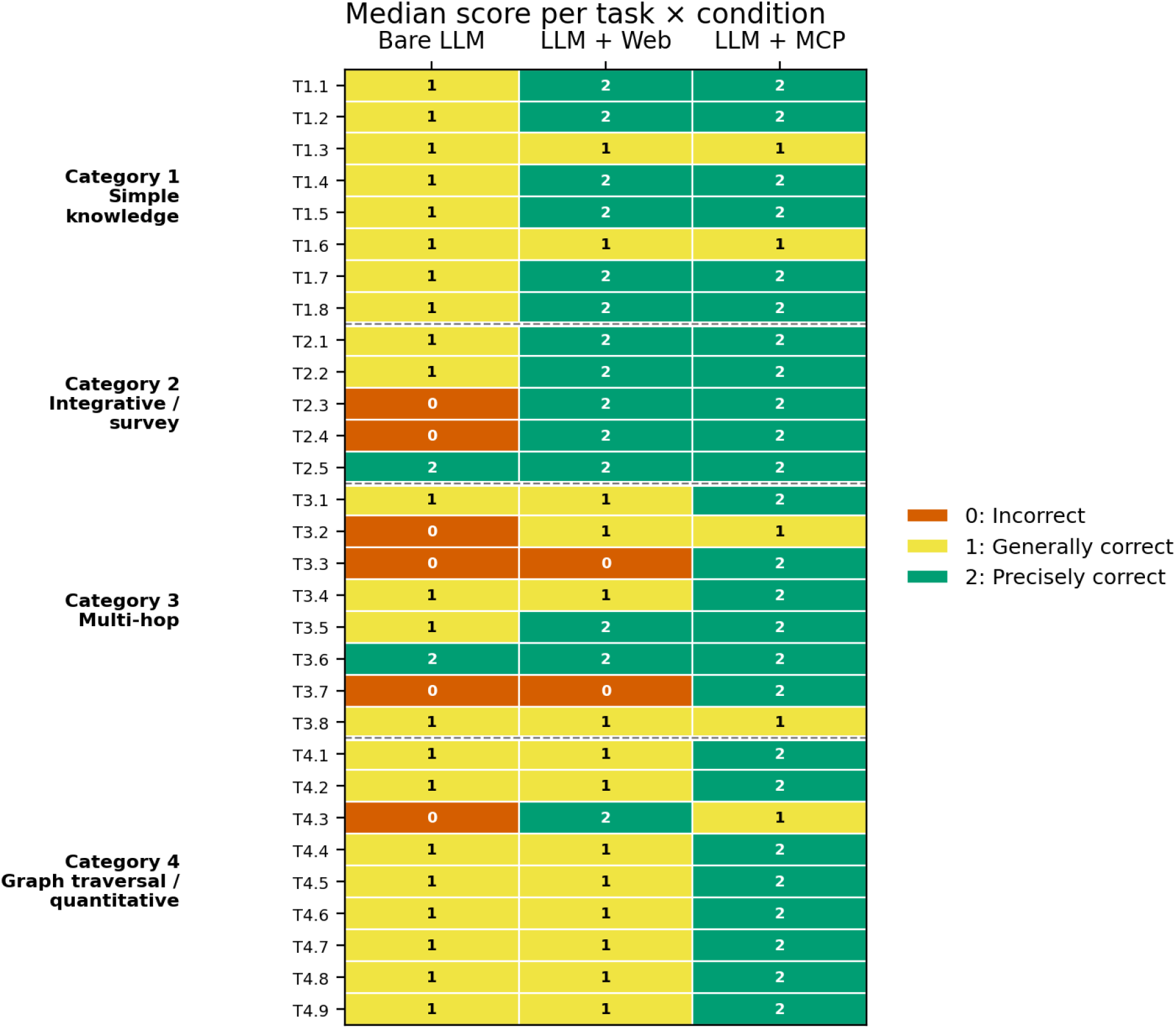
Score per task (rows) × condition (columns), grouped by category: Cell colour encodes the score (0/vermillion: incorrect, 1/yellow: generally correct, 2/green: precisely correct). MCP scores 2 across every category with a small number of exceptions. Web performs similarly to MCP within categories 1 and 2, but responses degrade in 3 and 4. Bare ceilings at 1 across most of the battery. For T3.3 and T3.7 Bare and Web both score 0 while MCP scores 2. No substantially fabricated results were produced.

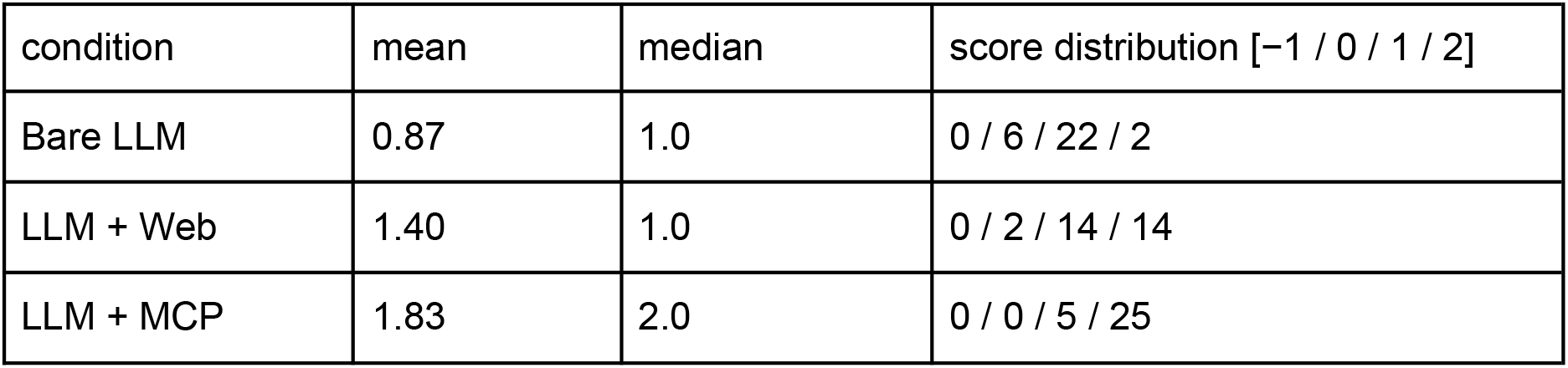

Pairwise differences across the three conditions were all highly significant (Friedman χ^2^(2) = 36.10, p = 1.4 × 10^−8^ for the omnibus; pairwise Wilcoxon signed-rank with Holm–Bonferroni correction: bare vs MCP p = 7 × 10^−6^, bare vs web p = 1.57×10^−3^, MCP vs web p = 3.00×10^−3^; n=30).

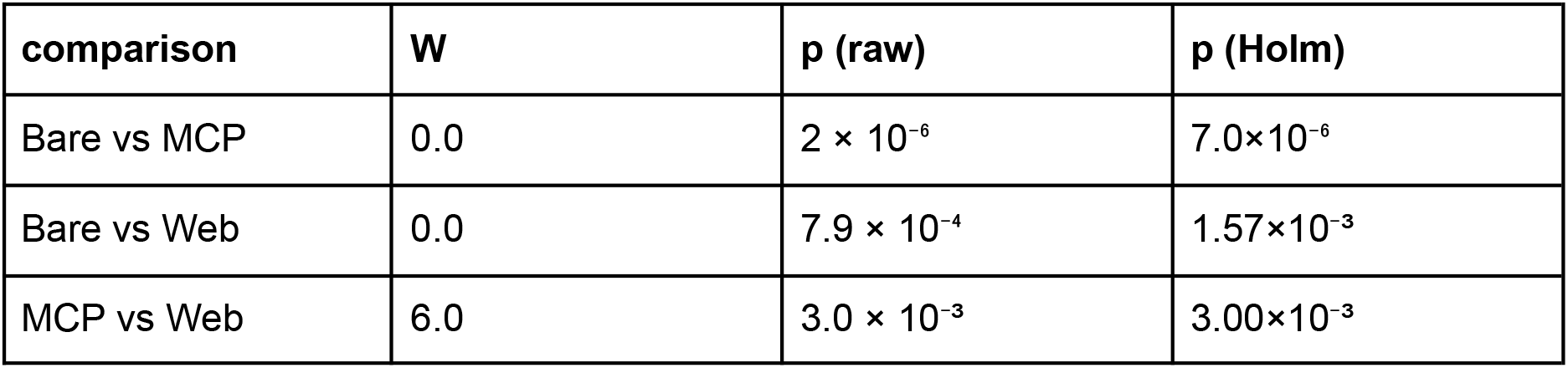

### The MCP advantage is greatest for multi-hop and quantification tasks

The fraction of tasks scoring “precisely correct” (score = 2) per category per condition (Figure 3), shows a higher differential between the MCP and web conditions as the categories require increasing quantification and integration across different data types. Per category performance:

**Figure 3.**
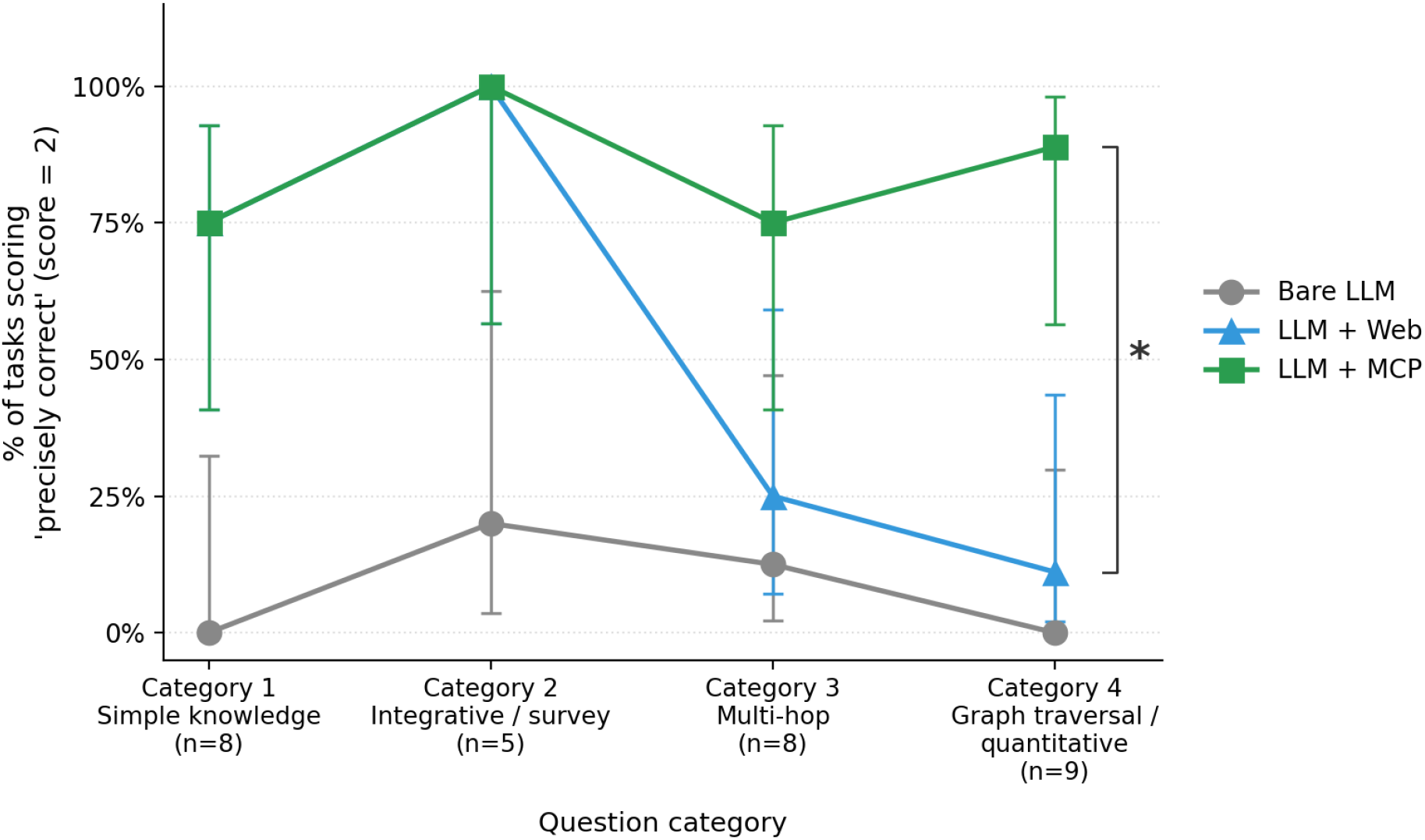
Rate of precise-correct answers (score = 2) per condition, by task category. Question categories against percentage of precisely correct answers per condition. Three lines: Bare LLM (grey), LLM + Web (blue), LLM + MCP (green) with 95% Wilson confidence intervals. Category 4 shows the greatest advantage for MCP, MCP 8/9 precise vs Web 1/9 vs Bare 0/9, where entity-precise quantitative answers are required. Category 1 (simple knowledge) and Category 2 (integrative survey) collapse the MCP/Web gap. At category 4, the MCP performed significantly better than the web condition W = 5.0, p = 0.0391, two-sided Wilcoxon signed-rank test.

– **Category 1 (simple knowledge, n = 8):** MCP and Web both achieve 75% precision (6/8). For well-documented knowledge, web search recovers the same answer as direct MCP access. Bare LLM scores 0/8 precise.
– **Category 2 (integrative / survey, n = 5):** MCP and Web both achieve 100% precision (5/5). On open-ended descriptive questions where the answer is enumeration of resources rather than entity-precise quantitation, web search again matches MCP. Bare LLM reaches 1/5.
– **Category 3 (multi-hop, n = 8):** MCP 75% (6/8) vs Web 25% (2/8) vs Bare 12% (1/8). MCP outperforms web as access to the VFB kb allows integration of datasets.
– **Category 4 (graph traversal / quantitative, n = 9): MCP 89% (8/9) vs Web 11% (1/9)**. The demands for quantification were achieved by the MCP condition, but the web condition could only do so if the exact data it needed was available in the literature. The bare condition failed to accurately quantify responses.

### Entirely or majority fabricated answers were not observed; however, partial fabrication did occur

No condition produced an answer scored as outright fabricated (score = −1) when read as a whole. The LLM used, Opus 4.7 within Claude Code, is reliable in flagging when the response is uncertain, and does not attempt a detailed answer when it does not have the information required. However, partial fabrication (PF flag: answer is generally correct but contains at least one fabricated detail) was observed at meaningful rates. The impact of partial fabrications on response usefulness depends not only on the rate, but also on whether the detail affected is important for the user. For example, a response accurately describing a circuit of interest can be made unreliable by a fabricated driver line identifier.

It is possible that the LLM does not flag the uncertainty of these partial fabrications if the model is confident in answering the question in general, but not some specifics. It is currently not possible to know why these errors occur, and the rate of occurrence, kind and severity of fabrications may vary from model to model. Therefore we reviewed and catalogued all partial fabrications for this LLM on our task battery and categorised them, reasoning that the broader kind of partial fabrications are more likely to be meaningful in the broader context than specific examples. From our review, we found that the rate of PF does not vary systematically with test condition (Figure 4), but the failure mode does:

**Figure 4.**
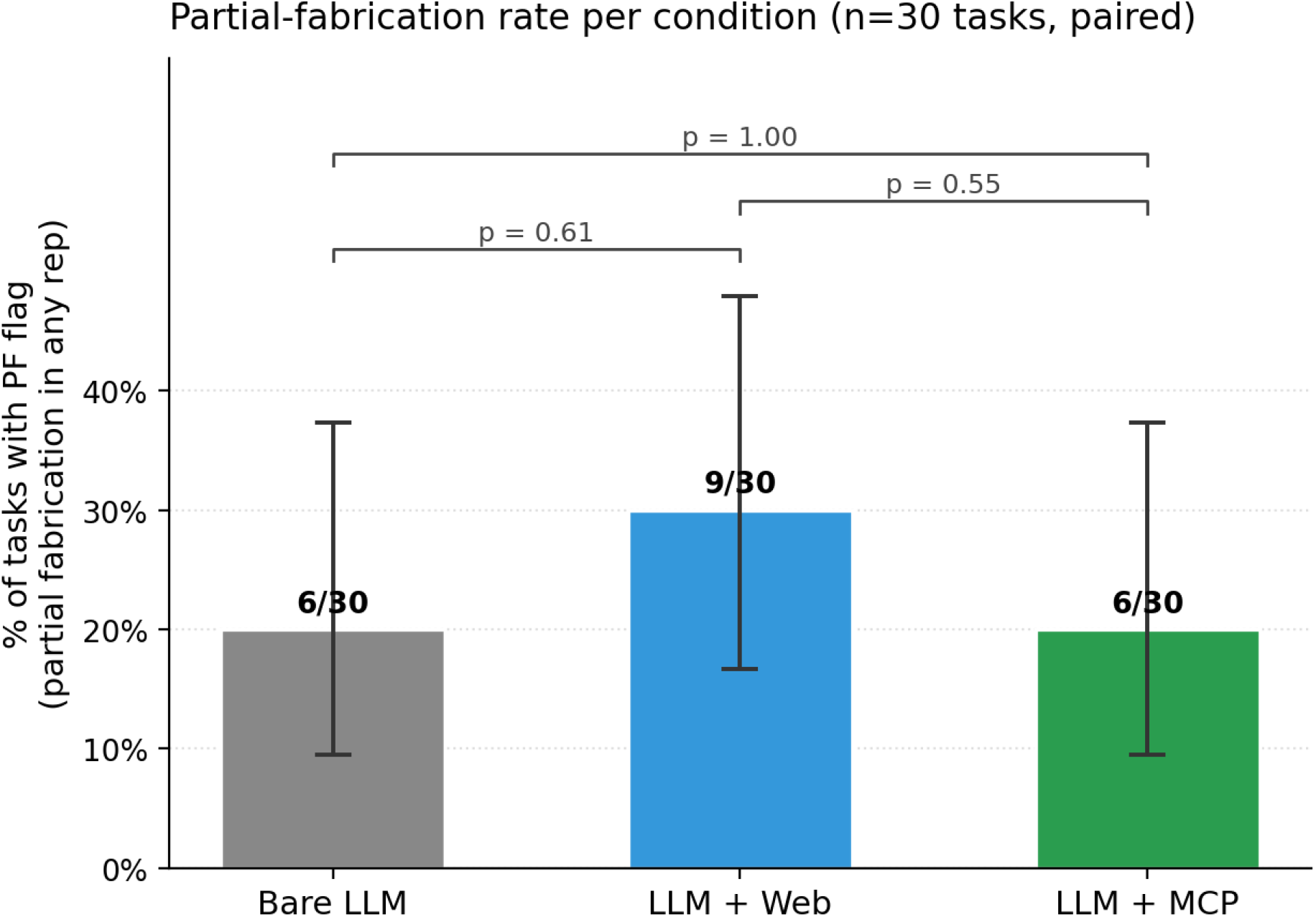
Partial-fabrication (PF) rate does not vary by condition: per-condition plot collated across all 30 tasks with pairwise McNemar tests on the paired binary outcomes. A task is counted as PF if any single partial fabrication was flagged. Overall PF rates are not statistically distinguishable across conditions (Bare 6/30, Web 9/30, MCP 6/30; all pairwise McNemar p > 0.5). 95% Wilson confidence intervals shown.

| Condition | PF rate (overall) | Identified failure modes |
| --- | --- | --- |
| Bare LLM | 6 / 30 (20%) | Cited paper does not contain the claim<br>Citation metadata wrong<br>Misattributed mechanism |
| LLM + Web | **9 / 30 (27%)** | Author transposition on real URL<br>Paper misattribution<br>Entity-binding within a real paper<br>Invention with citation<br>Wrong identifier<br>Wrong factual claim |
| LLM + MCP | 6 / 30 (20%) | Training-data fill on top of correct MCP scaffold<br>Ontology-class drift / confusion<br>Aggregation level wrong<br>Invented small claim with no MCP scaffold grounding |

### Failure-modes vary by condition

For the bare-LLM condition, three subclasses of errors were identified from eight partial fabrications occurring in the six tasks with the PF flag:

| Subclass | Tasks | Pattern |
| --- | --- | --- |
| Cited paper does not contain the claim | T2.2, T2.4, T3.1, T3.3 | Paper named is real, claim attributed to it is not there |
| Citation metadata wrong | T3.6, T4.3 | Paper is real but year/author list/journal misremembered |
| Misattributed mechanism | T3.6, T2.4 | Right family of fact, wrong specific entity |

Without any additional context to draw on, the LLM can only provide general responses to questions from its training data. The failure-modes observed suggest that this lack of context, which prevents the LLM from fact checking against papers, causes errors in citations where the claim is not supported at all, or the citation metadata is wrong, or the factual information is confused and attributed to the wrong specific entity. Our prompting to not report facts without references likely reduced the occurrence of further factual claims which are more poorly represented in the training data. This result confirms that LLMs without additional context may fail to provide claims supported by the literature.

The LLM + web condition aims to improve on the bare-LLM by gathering relevant context, primarily from research and review papers, to overcome the issue of ‘misremembering’ literature facts by including a summary of those papers in its context window. However, this causes a separate set of failure-modes:

| Subclass | Tasks | Pattern |
| --- | --- | --- |
| Author transposition on real URL | T1.7, T1.8, T2.5, T4.4, T4.7 | URL is to correct paper; first author misnamed in answer |
| Paper misattribution | T2.3 | Correct content, wrong source paper entirely |
| Entity-binding within a real paper | T4.1, T4.4 | Connection real, total right, but spatial/directional binding swapped |
| Invention with citation | T4.4 , T4.8 | Claim not in cited source at all |
| Wrong identifier | T4.6 | Real identifier system, wrong specific value |
| Wrong factual claim | T4.7 | No citation problem; just wrong biology |

These errors mostly fall under either a repeated “real paper, real URL, but they do not match” pattern or a “real paper, correct context, but misunderstood” pattern. The LLM can still get confused by complex papers and misinterpret them.

Under the LLM+MCP condition the most striking change to the failure-modes is the lack of errors in facts drawn from the VFB kb. Instead errors arise from adding data on top of the ground-truth scaffold from the MCP tool calls, or from misinterpreting the correct level of ontology class to use:

| Subclass | Tasks | Pattern |
| --- | --- | --- |
| Training-data fill on top of correct MCP scaffold | T4.2 | MCP returned right ID; model added wrong metadata from memory |
| Ontology-class drift / confusion | T3.3, T3.5 | Started right, conflated sibling/parent classes mid-answer |
| Aggregation level wrong | T4.1, T4.7 | Correct entities and direction, wrong level of resolution |
| Invented small claim | T1.2 | Free-form claim with no MCP grounding |

These classes of errors can be minimised further by 1) allowing MCP tool access to papers and 2) providing further guidance in how to use the data returned by the VFB-MCP tool calls (see discussion).

## Discussion

### VFB-MCP makes *Drosophila* neuroscience data accessible

Our results show that VFB-MCP enables accurate, sourced and quantified responses to *Drosophila* neuroscience questions. This opens up *Drosophila* neuroscience data to researchers without deep knowledge of the field, and/or the time and coding knowledge to use APIs. This work builds on previous work showing that an MCP layer over a curated biological resource substantially improves LLM accuracy for relevant questions.^19,22^ We also show the effectiveness of LLM introspection on MCP responses over multiple turns to deeply reason over and acquire context from ontology backed, expert-curated knowledgebases. This allows the LLM to access diverse data curated for specific entities, and understand the connections between them, which is vital for accessing the expanding connectomics datasets available for *Drosophila*. Future connectomes will be greater in scope and complexity and will likely also benefit from this approach to accessing them in the further context of the model organism’s neurobiology.

### Entity binding is a core advantage of structured tool access over text retrieval

The failure modes catalogued in the results section all share a structural property: text-retrieval pipelines (traditional RAG, web search, LLM training-data recall) cannot enforce that the entity to which the LLM attaches quantified results is the entity the source actually describes. This is shown with the web condition incorrect entity binding and incorrect identifier errors, whereas the MCP condition reliably attached quantitative data to the correct entity. This failure is not eliminated by frontier LLMs (Opus 4.7); it is a property of the LLM inferring entity binding from text. The MCP-mediated alternative for the same questions runs along a different track: a `run_query(id, query_type)` call is keyed on the entity ID, so misbinding is impossible by construction. The value returned for entity A is, by the design of the protocol, about entity A.

### Advantages and disadvantages of the web and MCP condition

Both web and MCP conditions are types of RAG. When using the VFB-MCP the VFB knowledgebase is the additional context, and when using web searches the information returned from web pages and papers are the context. The first gap between these methods is the available data. Large datasets are produced for *Drosophila* neuroscience at an increasing rate, and text from papers alone cannot include all of the information from these datasets. Therefore if there is no paper that happens to contain the quantitative, or qualitative data the user is searching for, then the web condition cannot find it. However, rhe MCP condition can make any query of a given dataset. Using connectomics as an example, papers only include a small fraction of all of the possible connectivity found in a full CNS connectome, while the VFB-MCP can return connectivity information for any given neuron in the dataset. This advantage is essential to the performance of the MCP condition. Compounded with the inherent disadvantage of attempting to extract entity binding information from text, the MCP condition has a substantial accuracy advantage for complex, quantitative questions.

A more subtle disadvantage for the web condition is that web fetches are actually a two-stage LLM pipeline when used within Claude Code, WebFetch passes each page through a separate summariser LLM and returns only its extraction, so any binding error or fabrication in a web answer could originate at either stage and is indistinguishable from outside the pipeline. The MCP condition avoids this requirement entirely, the structured data returned can be logically searched directly. However, MCP does not eliminate all hallucinations: The T4.2 (olfactory pathway) failure mode shows that even when the architectural scaffold of an answer is correctly grounded in MCP queries, the LLM can stack training-data-recall on top with citations and quantitative details that have no MCP basis. Importantly, the failure-mode is qualitatively quite different: with MCP, the scaffold is reliable and the failure occurs in adding in data from other sources; with web, the entire response is built from synthesising multiple sources any of which may be misinterpreted. As these two conditions have differing failure-modes, it may be possible to supplement the MCP with web searches to build on the strengths of both methods.

### Agent-mediated verification of responses is slow and costly

A separate LLM agent can be spawned after each response to fact-check each quantitative claim against its cited sources before the answer is shown to the user. We used this approach ad hoc during scoring when the human reviewer was unable to find evidence for claims made in a response, as a secondary check. Three caveats keep it from being integrated into the VFB-MCP condition. First, it is slow: each per-claim verification call takes tens of seconds (search, fetch source, read, render verdict). Multiplied by the number of claims in an answer this can double or triple latency and cost. Second, the verifying agent has the same vulnerabilities as the initial model did. Third, verification requires further token usage. We therefore did not use verification agents in this assessment, although potentially using a lighter model to confirm IDs and papers exist could reduce certain errors without significant extra cost.

### MCP over direct API access for LLMs

An alternative to the MCP server would be to simply use the LLM to write code against the existing programmatic API, vfb_connect (see data and code availability). We see three reasons to prefer MCP. First, the LLM selects a tool and fills in typed parameters rather than writing arbitrary code; this eliminates an entire class of errors (syntax, import, authentication, response parsing). Second, MCP tools encode domain expertise in their semantics: an operation like `run_query(id, “NeuronInputsTo”`) is a research-level abstraction over the underlying Cypher query, whereas using the API requires the LLM to know method names, parameter conventions, and how to chain calls. Third, any LLM with MCP compatibility can be used without changes to the server.

### Limitations

The evaluation of VFB-MCP presented here has several limitations. (i) The evaluation is restricted to Opus 4.7, while the VFB-MCP can in principle be used by any capable LLM. (ii) We performed 1 rep per task and therefore the impact of the probabilistic nature of LLM responses is not captured here. (iii) The task battery, while designed to span representative categories, does not exhaust the space of possible neuroscience queries. Further, web and bare condition tests are highly dependent on whether there are publications with information for the given cell classes. For well-studied neurons, even connectomic quantitative data can be found in papers, for less-studied neurons these conditions will have no data in papers to work with. In contrast, as the data held in the VFB kb is now dense, connectivity for any given neuron can be found. (iv) The assessor was not blinded to condition as the provenance citations are inherently distinguishable between conditions. (v) The VFB-MCP server’s `run_query` can return very large payloads (1 MB+ for broad connectivity queries) which the harness must handle. (vi) The training-data-fill failure mode on VFB-MCP LLM responses motivates a planned literature-MCP enhancement to add a PubMed MCP server alongside VFB-MCP, which will require further testing.

### Minimising cost

Other than functional limitations, cost is perhaps the most important factor in VFB-MCP use. For our assessment of Opus 4.7 against the task battery, we limited the cost per prompt to $3 USD. Because the VFB-MCP condition cost 12x the bare condition, this limit was included in the LLM context and reduced the number of tool calls made. For users exploring data with multiple LLM queries in a session this cost could quickly become substantial. Options for limiting cost are primarily to only use tool calls when necessary and to use cheaper models, for which the use of MCP tools has been shown to close the response quality gap with frontier models.^22^

Another option is for the resource to take on the cost of running the model. We are developing a VFB-chat which also interfaces with the VFB knowledgebase but runs on a local model: Llama `meta-llama/Llama-3.3-70B-Instruct` hosted on Edinburgh ELM at https://elm.edina.ac.uk/api/v1. Local models are open and not subject to token costs. However, they do, incur energy and infrastructure costs, and require careful harnessing to ensure questions are relevant and to prevent potential abuse. We plan to make VFB-chat available to researchers to further increase the accessibility of the VFB knowledgebase.

### VFB-MCP does not present an increased security risk

The three VFB-MCP tools are predefined, read-only operations against a public knowledge resource that is already exposed via REST and Python APIs. There are no write paths and no arbitrary-code-execution paths in the MCP surface; an LLM client cannot invent a tool that does not exist on the server, cannot pass query strings interpreted as code, and cannot affect server state. The security threat is no greater than that from the existing public-API.

### Skills as a complementary mechanism

MCP servers define what tools exist. For VFB-MCP, the LLM is given a set of tools and the information on how to use them to query the knowledgebase. A complementary mechanism emerging on the model side is the skills abstraction (e.g. Claude Skills): packaged procedural knowledge including natural-language instructions, code snippets and worked examples, that the model can load on demand for tasks requiring more than a single tool-call sequence. Skills therefore provide context to inform an LLM of a tested pattern of how to accomplish a task with the available tools.^32^ For exploratory biological work that combines multiple VFB-MCP tool calls, literature retrieval, and code, skills could pre-package successful tool-call patterns from prior research workflows, lowering the threshold for less-experienced users to reach expert-grade outputs. The MCP provides the tool surface and skills provide expert curated workflows using them. We anticipate that mature biological MCP ecosystems will be paired with curated skill libraries that encode community best-practice for common research workflows.

### An MCP ecosystem for biology

VFB-MCP is one node in a rapidly growing network of biological MCP servers. As this ecosystem matures, the full potential of MCP will lie in allowing an LLM to simultaneously query VFB for *Drosophila* circuit structure, OmniPath for molecular interactions,^1^ STRING for protein networks,^33^ UniProt for protein function,^34^ and PubMed or ASTA-MCP for literature,^35^ bringing together results from varied resources in a single conversational workflow. The standardised MCP protocol makes this interoperability possible with minimal work from the resources and the user. We expect that ontology-backed knowledge graphs, with their native support for relationship traversal and semantic reasoning, will serve as nodes in this network that facilitate access to deep, structured knowledge. Data from the knowledgebases remain as the ground truth scaffold, and the information from other sources can build on this. Ensuring confabulation does not increase will require careful harnessing. Registries such as BioContextAI may serve as centralised hubs for identifying relevant MCP servers.^36^

## Resource availability

### Contact

Requests for further information or resources should be directed to the lead contact, J. Douglas Armstrong or Alex D. McLachlan.

### Materials availability

This study did not generate any new, unique reagents.

### Data and code availability

– All data and code required to reproduce the benchmark results, statistics, and figures are available at https://github.com/VirtualFlyBrain/vfb-mcp-paper-supplement and archived at Zenodo (https://doi.org/10.5281/zenodo.21143107), under a Creative Commons Attribution 4.0 International (CC BY 4.0) licence. This archive contains the 30-task battery, the per-task ground-truth mark schemes, the complete set of expert scoring sheets for all three conditions (bare, LLM + web, LLM + MCP), the parsed scores and computed statistics, the benchmark runner, source data for Figures 2, 3, and 4 and the analysis and figure-generation scripts.
– The VFB-MCP server code is available at https://github.com/VirtualFlyBrain/VFB3-MCP under the MIT licence; the server is deployed at https://vfb3-mcp.virtualflybrain.org.
– The Drosophila Anatomy Ontology is available at purl.obolibrary.org/obo/fbbt.owl under CC BY 4.0.
– The vfb_connect Python API is available via PyPI.

## Acknowledgments

This work was supported by the Wellcome Trust grant 223741/Z/21/Z.

## Author contributions

Conceptualisation, R.C., J.D.A., A.D.M.; methodology, R.C., J.D.A., A.D.M.; investigation, A.D.M., R.C., C.P; development, R.C.; funding acquisition, D.O-S., J.D.A., G.S.X.E.J., N.H.B.; writing – original draft, A.D.M.; writing – review & editing, J.D.A., K.L; supervision, J.D.A., R.C.

## Declaration of interests

The authors declare no competing interests.

## Methods

### METHOD DETAILS

#### VFB-MCP server implementation

The VFB-MCP server is implemented in TypeScript (v5.9.3) using the MCP SDK (v1.26.0) with Express.js for HTTP transport. The server translates MCP tool calls into queries against VFB backend services. The server implements the HTTP transport variant of the MCP specification, operating as a stateless JSON-over-HTTP service. It is deployed as a Docker container (Node.js 18 Alpine base) with resource limits, running as a non-root user with a read-only filesystem. All benchmarks reported here were run against VFB-MCP server v1.8.1 at https://vfb3-mcp.virtualflybrain.org and v1.9.0 of VFB-Query.

The deployed server exposes **three generic tools**: `search_terms, get_term_info, run_query` combined with per-entity query introspection. `get_term_info` returns, for each entity, a Queries array enumerating the valid `query_type` values applicable to that entity. Available query types vary per entity; clients are directed by the server to call `get_term_info` first to enumerate them.

#### Example MCP Tool use flow

Our implementation uses a **search → introspect → execute** pattern. The ontology drives discovery at runtime rather than the tool list encoding it statically. In the cases of an LLM asked *“what neuron types provide input to the MBON-γ1pedc>α/β neuron, and which are dopaminergic?”* The model’s first call,

`search_terms(query=“MBON-gamma1pedc>a/b”)`, returns the canonical VFB ID (`FBbt_00100246`). Its second call, `get_term_info(id=“FBbt_00100246”)`, returns the entity’s classification, metadata, *and* a `Queries` array listing query types valid for this neuron, including `ListAllAvailableImages, PartsOf, SubclassesOf, ExpressionOverlapsHere, TransgeneExpressionHere, DownstreamClassConnectivity and UpstreamClassConnectivity`. Had the entity been, for example, a brain region, `get_term_info` would have returned a different Queries set (e.g. `NeuronsPartHere, ExpressionOverlapsHere`). The model selects `UpstreamClassConnectivity` and issues `run_query(id=“FBbt_00100246”, query_type=“UpstreamClassConnectivity”)`, receiving the structured list of upstream neuron classes with synapse weights, which it can then filter for dopaminergic types via a follow-up `search_terms` call with the appropriate facet. Three generic tools and one ontology-driven introspection step suffice to answer a question that, in a static-tool architecture, would require either a bespoke `query_inputs_to_mbon` tool or LLM-generated graph-database queries against an exposed schema.

#### Neo4j graph database and ontology layer

VFB’s knowledge is stored in a Neo4j graph database where nodes represent biological entities (neurons, brain regions, cell types, genes, transgenes, publications) and edges represent typed relationships (synapsed_to, part_of, subclass_of, expresses, has_image, etc.). The Drosophila Anatomy Ontology (DAO) provides the semantic backbone, ensuring that entities are classified within a formal hierarchy and that relationships between them are semantically constrained.

Graph traversal operations (connectivity queries, hierarchical navigation, morphological similarity searches) are implemented as parameterised Cypher queries executed against Neo4j. Each query type listed in the per-entity `Queries` array maps to a specific Cypher template; `run_query` resolves the `(id, query_type)` pair to the appropriate template and parameters. This native graph traversal is substantially more efficient and expressive than equivalent operations over relational databases, which would require recursive CTEs or application-level iteration.

#### Benchmark task battery design

We designed 30 neuroscience tasks across four categories (see results for the categories and data and code availability for the complete battery). Tasks were designed by the VFB team, to reflect questions that arise in real research workflows, framed broadly enough that a bare LLM could attempt an answer from training data but precisely enough that database-grounded answers are verifiably more specific. Tasks were designed so that data from VFB could be used to ground the answers, but could also use web-retrieved context or training data. Specifically, no question explicitly requested data from the VFB kb.

#### Benchmark conditions and runner architecture

The benchmark compares three conditions:

1. **Bare LLM**: no tools of any kind. The LLM answers from training data only.
2. **LLM + VFB-MCP**: an agentic configuration where the LLM has access to the three VFB-MCP tools described above and nothing else.
3. **LLM + web**: a second agentic configuration where the LLM has access to web search and web fetch tools (`WebSearch, WebFetch` in Claude Code) but no MCP. Importantly, the primary LLM never sees raw page content in this condition: `WebFetch` takes a URL plus an extraction prompt; the harness fetches the page, runs a separate (typically smaller) model over the page content with that prompt, and returns only the summary or extraction. The +web condition is therefore a two-stage LLM pipeline: page → summariser LLM → summary → primary LLM → answer. `WebSearch` similarly returns ranked summary snippets rather than full pages.

We benchmarked one LLM across the three primary conditions: **Claude Opus 4.7**.

**Opus runner**: questions were dispatched via the Claude Code CLI in non-interactive mode (`claude –p`), with `--output-format stream-json –verbose --no-session-persistence --strict-mcp-config`. Per condition, the runner either (a)supplied an empty MCP configuration with `--tools ““` to disable all tools (bare), (b) supplied the VFB-MCP server configuration with `--allowed-tools` listing the VFB-MCP tools (MCP arm), or (c) supplied an empty MCP configuration with `--tools “WebSearch” “WebFetch”` to expose web tools only (web arm). A `--max-budget-usd 3.00` cap and a 600 s wall-clock subprocess timeout protected against runaway loops.

#### Per-call telemetry

Each repetition records (full schema in the released codebase):

– The full sequence of tool calls (`tool_trace`): name, id, input arguments, the corresponding tool result (truncated at 4 kB), an `is_error` flag.
– Derived call-category counts: `vfb_calls` (calls to `mcp virtual-fly-brain *), web_calls (WebSearch, WebFetch), vfb_web_calls` (the subset of `WebFetch` calls whose URL is on virtualflybrain.org), infra_calls (built-in tools used for filesystem-style bookkeeping), `cli_calls` (CLI deferred-tool-schema fetches).
– Token usage and cost: `input_tokens, output_tokens, cache_read_input_tokens, cache_creation_input_tokens`, and `cost_usd` (taken from the API’s `total_cost_usd`, billing-grade metering from the API response envelope, not a model-side estimate).
– Timing: `duration_ms, duration_api_ms, wall_time_ms`.
– Final-event metadata: number of model turns, success flag, exit code on failure.
– Per-rep `model_alias` (the alias passed) and `resolved_model` (the model identifier the CLI’s stream-json init event reported), so the resolved model is recoverable from per-rep records.

#### Oversized MCP responses and CLI spillover

For some queries, particularly `run_query` with broad connectivity types on heavily-connected nodes, the VFB-MCP server returns very large JSON payloads (single calls observed up to 1.2 MB and 40,000+ rows). The Claude Code CLI handles such oversized tool responses by spilling them to a session log file on disk and returning a short text indicating the file path. The model then uses built-in `Read/Grep/Bash` tools to parse the spilled content. For a typical heavy connectivity question on Opus this can inflate apparent tool-call counts by 10–20 rounds, all of which are file-handling rather than domain queries. We therefore treat `vfb_calls` as the primary tool-use count, and treat `infra_calls` as a secondary measurement of CLI-implementation overhead. Therefore, to use the VFB-MCP it is highly recommended that you use a harness that can handle large response payloads.

#### Observed cost and runtime

A single 1-rep sweep of all 30 tasks on Opus 4.7 cost approximately: Bare arm $1.36 (~14 min wall-clock); LLM + VFB-MCP arm $16.30 (~61 min); LLM + web arm $11.80 (~43 min). Cost ratios in the 1-rep sweep are MCP/bare ≈ 12×, web/bare ≈ 9×, MCP/web ≈ 1.4×.

#### Condition-specific system prompts

We used three condition-specific prompts:

– **BARE_SYSTEM_PROMPT**: declarative (“You do NOT have access to any tools, databases, or the web — you cannot query [enumerated systems]”), explicit prohibition on claiming tool use, explicit prohibition on inventing identifiers, constrained source vocabulary (“either ‘general training knowledge’ or a specific publication you are confident about”).
– **WEB_SYSTEM_PROMPT**: declarative (“You have access to web search and web fetch tools and you MUST use them”), tool-use mandate, with citations requiring the specific URL fetched or search query run.
– **MCP_SYSTEM_PROMPT**: declarative (“You have access to Virtual Fly Brain database tools and you MUST use them”), tool-use mandate, per-call provenance citation requirement.

#### Scoring protocol

All answers from each condition were scored by a domain expert. VFB’s current knowledgebase state served as the primary ground truth. Because the LLM was prompted to reference every claim, the claim must be supported by the referenced paper in each case except for very general knowledge claims. If the claim was not covered by the VFB knowledgebase then the referenced paper was deemed sufficient.

Answers were scored on a 4-point scale:

– **−1 (Fabricated)**: Answer is substantially fabricated and dominated by false specifics. The central facts are incorrect and not supported by the literature. (invented neuron types, non-existent driver lines, fabricated citations, made-up synapse counts).
– **0 (Incorrect)**: Wrong answer from genuine reasoning errors or lack of knowledge, not fabrication.

To differentiate −1 and 0, consider an answer to a question asking for Kenyon cell subtypes. If the response claims that Kenyon cells have subtypes but it cannot provide that information, that would be scored as 0. If the model confabulated subtypes that do not exist, that would be scored as −1.

– **1 (Generally correct)**: Qualitatively right, consistent with published literature, correct in broad strokes, but lacks specificity.
– **2 (Precisely correct)**: Specific, quantitative, and verifiable against current VFB data (or literature), includes correct identifiers, synapse counts, or ontological relationships from the database.

The primary differentiator between 1 and 2 is the presence of quantitative information where appropriate.

Two independent flags were recorded:

– **PF (Partial fabrication)**: Answer scores 1 or 2 overall but contains at least one specific fabricated detail (e.g., a non-existent citation or invented driver line name peripheral to the main answer).
– **LD (Literature-data discrepancy)**: Published literature and current VFB data disagree on this question. A property of the question, identified during ground-truth verification. In LD cases, a bare/web answer correctly reporting the literature consensus scores 1 (relative to its source), while an MCP answer correctly reporting current database state scores 2 (relative to the primary ground truth). Neither is penalised for the discrepancy itself.

The assessor was not blinded to condition: the provenance citations in MCP answers (referencing specific VFB queries) and the URL citations in web answers are inherently distinguishable from bare-LLM reliance on training knowledge, making effective blinding impractical.

### QUANTIFICATION AND STATISTICAL ANALYSIS

Task scores were used as the unit of analysis (n = 30 tasks, 3 conditions: bare LLM, LLM + VFB-MCP, LLM + web). We tested the global null that all three conditions share an identical distribution with Friedman’s test (χ^2^(2) = 36.10, p = 1.4 × 10^−8^). Given a significant omnibus, we conducted pairwise Wilcoxon signed-rank post-hoc tests with Holm–Bonferroni correction reported as the primary family-wise error control; Bonferroni and Benjamini–Hochberg false-discovery-rate corrections concur. All three pairwise comparisons rejected the per-pair null at α = 0.05 under all three corrections. For the bare-vs-MCP comparison the Wilcoxon statistic was W = 0, reflecting the fact that MCP equalled or exceeded bare on every task in the battery (25 MCP > bare, 5 ties, 0 bare > MCP); scipy uses the normal approximation rather than the exact distribution when ties are present. Analysis code (`analyse_results.py`) is available in the supplementary material.

For the per category statistics, we conducted one pre-specified post-hoc Wilcoxon signed-rank test: MCP versus Web on Category 4 (graph traversal / quantitative) (Wilcoxon W = 5.0, p = 0.039). This was chosen prior to per-category inspection on the grounds that the architectural hypothesis is strongest where entity-precise quantification is required. Per-category proportions in Figure 3 (with Wilson 95% CIs) are otherwise descriptive.

## Notes

https://doi.org/10.5281/zenodo.21143107

### Competing Interest Statement

The authors have declared no competing interest.

### Summary of Updates

Version 2 changes: - Supplementary data and code is now available in the linked Zenodo archive and GitHub repo. Updated the Data and Code Availability statement accordingly. - Refined Figures 3 and 4 (consistent condition ordering and colour scheme; exact p-values shown). - Results and conclusions are unchanged.

